# Cellular mechanotransduction of human osteoblasts in microgravity

**DOI:** 10.1101/2024.03.03.583164

**Authors:** Nadab H. Wubshet, Grace Cai, Samuel J. Chen, Molly Sullivan, Mark Reeves, David Mays, Morgan Harrison, Paul Varnado, Benjamin Yang, Esmeralda Arreguin-Martinez, Yunjia Qu, Shan-Shan Lin, Pamela Duran, Carlos Aguilar, Shelby Giza, Twyman Clements, Allen P. Liu

**Author notes:** Corresponding authors: Twyman Clements, Allen P. Liu. Equal contribution (Nadab H. Wubshet, Grace Cai).

## Abstract

Astronauts experience significant and rapid bone loss as a result of an extended stay in space, making the International Space Station (ISS) the perfect laboratory for studying osteoporosis due to the accelerated nature of bone loss on the ISS. This prompts the question, how does the lack of load due to zero-gravity propagate to bone-forming cells, human fetal osteoblasts (hFOBs), altering their maturation to mineralization? Here, we aim to study the mechanotransduction mechanisms by which bone loss occurs in microgravity. Two automated experiments, 4 microfluidic chips capable of measuring single-cell mechanics of hFOBs via aspiration and cell spheroids incubated in pressure-controlled chambers, were each integrated into a CubeLab deployed to the ISS National Laboratory. For the first experiment, we report protrusion measurements of aspirated cells after exposure to microgravity at the ISS and compare these results to ground control conducted inside the CubeLab. Our analysis revealed slightly elongated protrusions for space samples compared to ground samples indicating softening of hFOB cells in microgravity. In the second experiment, we encapsulated osteoblast spheroids in collagen gel and incubated the samples in pressure-controlled chambers. We found that microgravity significantly reduced filamentous actin levels in the hFOB spheroids. When subjected to pressure, the spheroids exhibited increased pSMAD1/5/9 expression, regardless of the microgravity condition. Moreover, microgravity reduced YAP expression, while pressure increased YAP levels, thus restoring YAP expression for spheroids in microgravity. Our study provides insights into the influence of microgravity on the mechanical properties of bone cells and the impact of compressive pressure on cell behavior and signaling in space.

## INTRODUCTION

Osteoporosis causes bones to become weak and brittle as individuals age and commonly leads to fractures with mild stresses or a fall. While dietary habits, hormone levels, and certain medical conditions are contributing factors in osteoporosis, it is also well appreciated that weight-bearing exercises are beneficial to the bones and lower the risk of osteoporosis. Physiological measurements during space flight have informed us that microgravity causes bone demineralization, skeletal muscle atrophy, vestibular problems, and reductions in plasma volume and red cell mass^1^. Many of these physiological changes also mirror changes in different pathophysiological conditions and diseases in humans on Earth. For instance, bone mineralization is seen during prolonged bed rest, although not to the same extent as in microgravity^1^. The well-known NASA twin astronaut study on the International Space Station (ISS) documenting various physiological and systems-level data over 25 months reveals bone breakdown and formation markers change over the duration of spaceflight^2^. The complex patterns are difficult to explain, and the cellular regulation of bone formation and breakdown in microgravity is even more enigmatic.

Osteoblasts are mature bone-forming cells and represent a specific stage in bone development. Within this population, a subset of osteoblasts undergoes further differentiation to become osteocytes. The role of osteoblasts is to deposit minerals and proteins to form the extracellular network when and where bone is most needed. This complex and dynamic process is tightly regulated by the opposing action of the osteoclasts (which resorb bone), the secretion of growth factors, and physical activity^3,4^. Changes in the balance between bone deterioration and formation by osteoclasts and osteoblasts, respectively, can cause either an increase in overall bone density or more commonly bone loss. The growth factors bone morphogenetic proteins (BMPs), which belong to the large transforming growth factor-β (TGF-β) family, are key regulators in bone formation. Genes activated by BMP via SMAD signaling pathways, such as RUNX2, osteocalcin, and alkaline phosphatase, are involved in the differentiation of mesenchymal stem cells to mature osteocytes^5,6^. With the emergent field of mechanobiology^7^, there is growing interest in understanding how physical forces affect bone development at the cellular level^4,8^.

Several studies have identified YAP/TAZ transcription factors as regulators of mechanosensitive cellular behaviors^9,10^, where high cell tension is associated with increased YAP nuclear translocation and transcriptional activity. Intriguingly, crosstalk exists between BMP signaling and YAP/TAZ nucleocytoplasmic shuttling, suggesting a convergence of mechanical and biochemical cues that may directly regulate osteoblast functions. YAP/TAZ has been shown to bind to pSMAD2/3 activated by TGF-β in human embryonic stem cells to regulate stem-cell self-renewal^11^, but the cytoplasmic function of YAP/TAZ on SMAD is controversial and may be cell type dependent^12,13^. YAP phosphorylation at S127 promotes the retention of YAP in the cytosol^14^, and thus it is plausible that cells under low tension may not respond to BMP due to the sequestration of SMAD1/5/9 by phosphorylated YAP. Our overall hypothesis is that reduced cell tension, due to the known disruption of the actin cytoskeleton in microgravity conditions, would lead to YAP cytosolic retention and attenuate BMP signaling in osteoblasts (**Figure 1**). Furthermore, we postulate that application of external stress to increase osteoblast cell tension would rescue their BMP signaling in microgravity.

**Figure 1.**
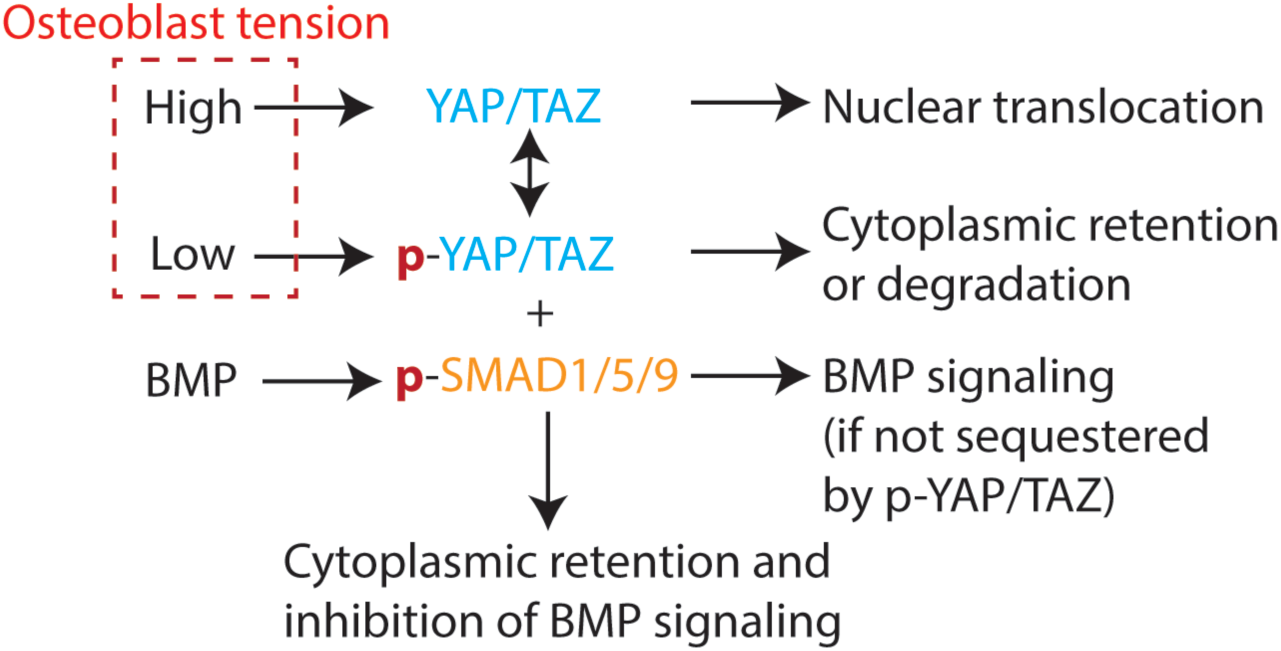
Hypothesis being examined in this study. High cell tension leads to YAP nuclear translocation whereas low cell tension leads to cytosolic YAP that sequesters phospho-SMAD proteins.

Herein, we describe the development of flight-ready hardware for two independent ISS experiments:1) microfluidic pipette aspiration for quantifying human osteoblast stiffness in microgravity and 2) stimulation of human osteoblast spheroids with and without applied pressure. The hardware was developed by Space Tango over multiple iterations of prototyping concepts with the key constraints of 12.30” by 8.15” by 4.26” for the CubeLab. The ISS experiments were flown on NG-18 on November 7, 2022, and on SpX-27 on March 14, 2023.

## RESULTS

### Hardware design for automated microfluidic pipette aspiration for the ISS

To characterize the mechanical properties of single cells, our lab has developed a polydimethylsiloxane (PDMS)-based microfluidics device that autonomously traps single cells and exerts known amounts of pressure based on varying fluid flow rates^15,16^. The working principle is based on micropipette aspiration, a well-established tool for single-cell mechanics research^17,18^. However, in conventional micropipette aspiration, a bulky setup with micromanipulation of a pulled glass micropipette is required. We have dramatically simplified the experimental setup to effectively a syringe pump to drive fluid flow through the PDMS device, thus enabling parallel measurements of multiple cells in a single chip. Cells are trapped in a series of trapping cups based on a hydrodynamic trapping scheme^16^. Instead of applying a negative pressure by moving a water reservoir in conventional micropipette aspiration, the microfluidic device aspirates on a trapped cell based on the pressure difference due to the pressure loss between the trapping cup and the end of the small channel that functions as a micropipette. The deformation of trapped cells as a function of increasing applied pressure via increasing flow rates can be used to determine their stiffness^16^. Per the requirements of the CubeLab, the PDMS chip was fabricated with constraints of a maximum of 8 mm chip height with the focus plane, which is the plane of the flow channel, not exceeding 4 mm height.

While the microfluidic device is relatively easy to operate on Earth, it still requires manual steps and the use of an epifluorescence microscope. For the experiment to be conducted on the ISS free of astronaut time (other than installing the CubeLab on the ISS), all the steps need to be automated and fit into the CubeLab. The required key capabilities are 1) automating cell lifting and resuspension, 2) keeping cells at 34 °C and other solutions in cold, 3) the ability to conduct experiments over multiple chips, and 4) brightfield and fluorescence imaging of cells in trapping cup structures.

Figure 2a illustrates the fluidic diagram. There are four cell culture bags maintained at 34 °C and cell lifting of each bag can be independently controlled. The setup also accommodates four PDMS chips, each paired with a cell culture bag. Cell culture media and phosphate buffer saline (PBS) are kept in the cold flask. A vibrating disc motor is attached to each of the four culture bags to create the agitation necessary to resuspend cells treated with TrypLE. To prevent bubbles from flowing in the PDMS chip, we employed a bubble trap. The fluidic manifold underneath the PDMS chips was machined to have inlet and outlet posts that match the inlets and outlets of the device. The PDMS chips were held in place by a compression plate with cutouts for the chips for brightfield and fluorescence imaging from the top (**Supplemental Figure 1**). The PDMS chips are to remain stationary while the 20x objective, illumination source, optics, and imaging sensors move in the *x* and *y* directions. The layout of the various components is shown in Figure 2b. The fully assembled hardware is shown in Figure 2c.

**Figure 2.**
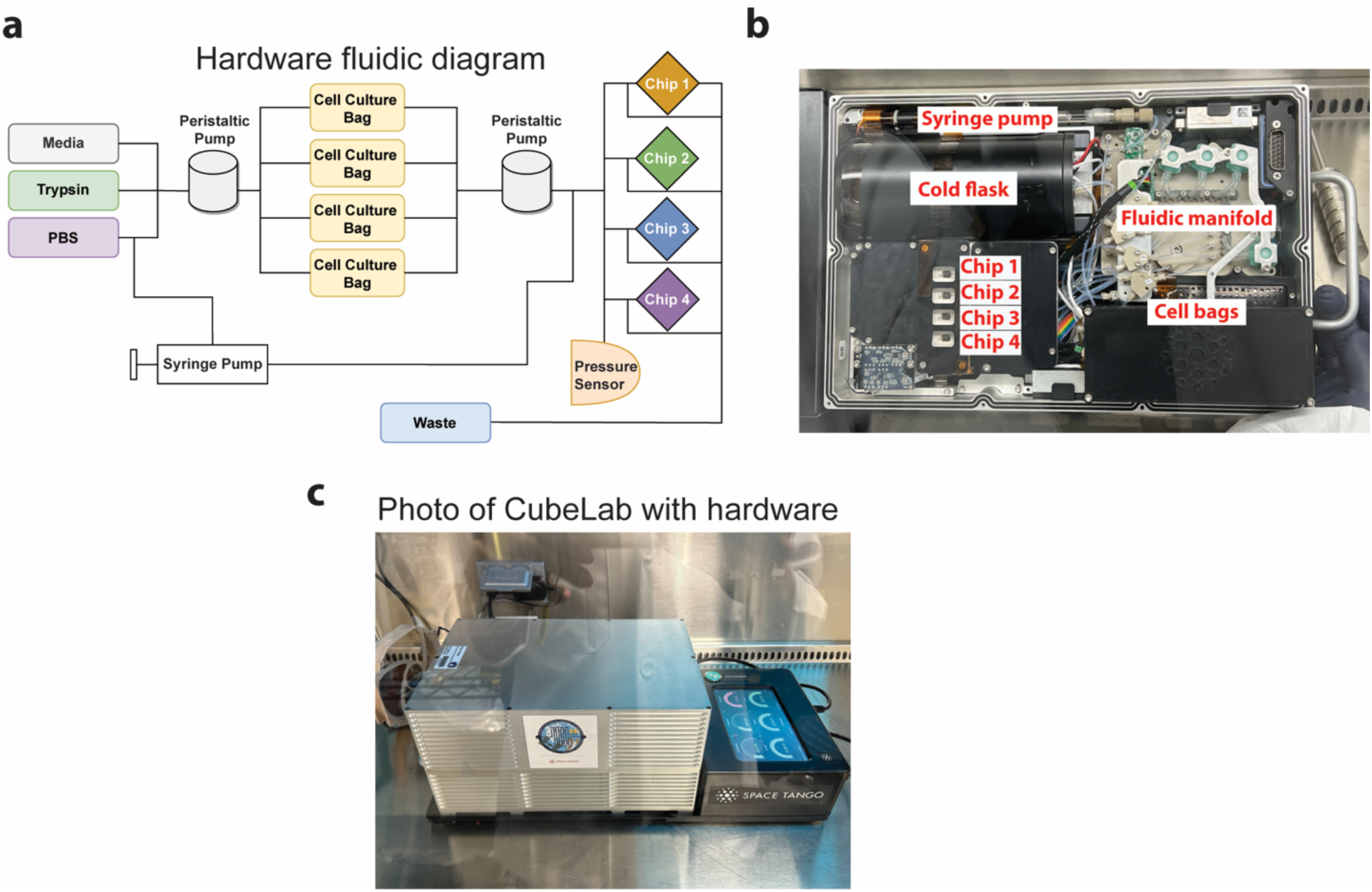
CubeLab for microfluidic pipette aspiration chip. **(a)** Schematic of Space Tango’s hardware fluidic diagram illustrates the fluidic routes and components of the integrated CubeLab. Pumps, valves, chips, and bag containers of cells, media, and waste are represented. **(b)** Inside of the CubeLab with major parts labeled. **(c)** Space Tango’s assembled CubeLab powered by an external source.

### Ground experiment using the microfluidics-integrated hardware

In our previous study, we have shown that the stiffness of human osteoblasts hFOBs is reduced when subjected to 6 hours of simulated microgravity using a random positioning machine^19^. This is consistent with the observed reduction in filamentous actin (F-actin) following simulated microgravity. Given cell tension is directly related to the density of cortical actin networks^20^, and that there is a positive relationship between cell stiffness and the amount of actin stress fibers^21^, we assert that cell stiffness directly relates to cell tension, and thus the relative cell tension can be inferred from our cell stiffness measurement.

We carried out a ground experiment using the CubeLab hardware that integrated the microfluidic chips with automated cell culture, cell lifting, and fluidic handling. Cells were labeled with Vybrant DiO in Kiyatec bags and allowed to grow for 2 days to simulate the duration of launch and ascent to the ISS. Media was then removed, and TrypLE was added to dissociate the cells. Agitation helped break up cell clumps into single-cell suspension and cells were loaded into the microfluidic chip at 0.2 psi. This is the pressure that allows fluid flow through our microfluidic chip and cells to be captured in the trapping cups. Increasing the flow rate by increasing the pressure of the syringe pump would increase the pressure exerted on the cell, as in micropipette aspiration (Figure 3a). Brightfield and fluorescence images were captured at each pressure increment of 0.1 psi from 0.2 to 2.0 psi (Figure 3b). We did not observe increased deformation with increasing pressure (Figure 3c), as we observed previously when using this microfluidic device^16,19^. Instead, the protrusion length into the microfluidic pipette channel remained constant throughout the experiment. It was not clear why the microfluidic device did not function as expected in the flight hardware.

**Figure 3.**
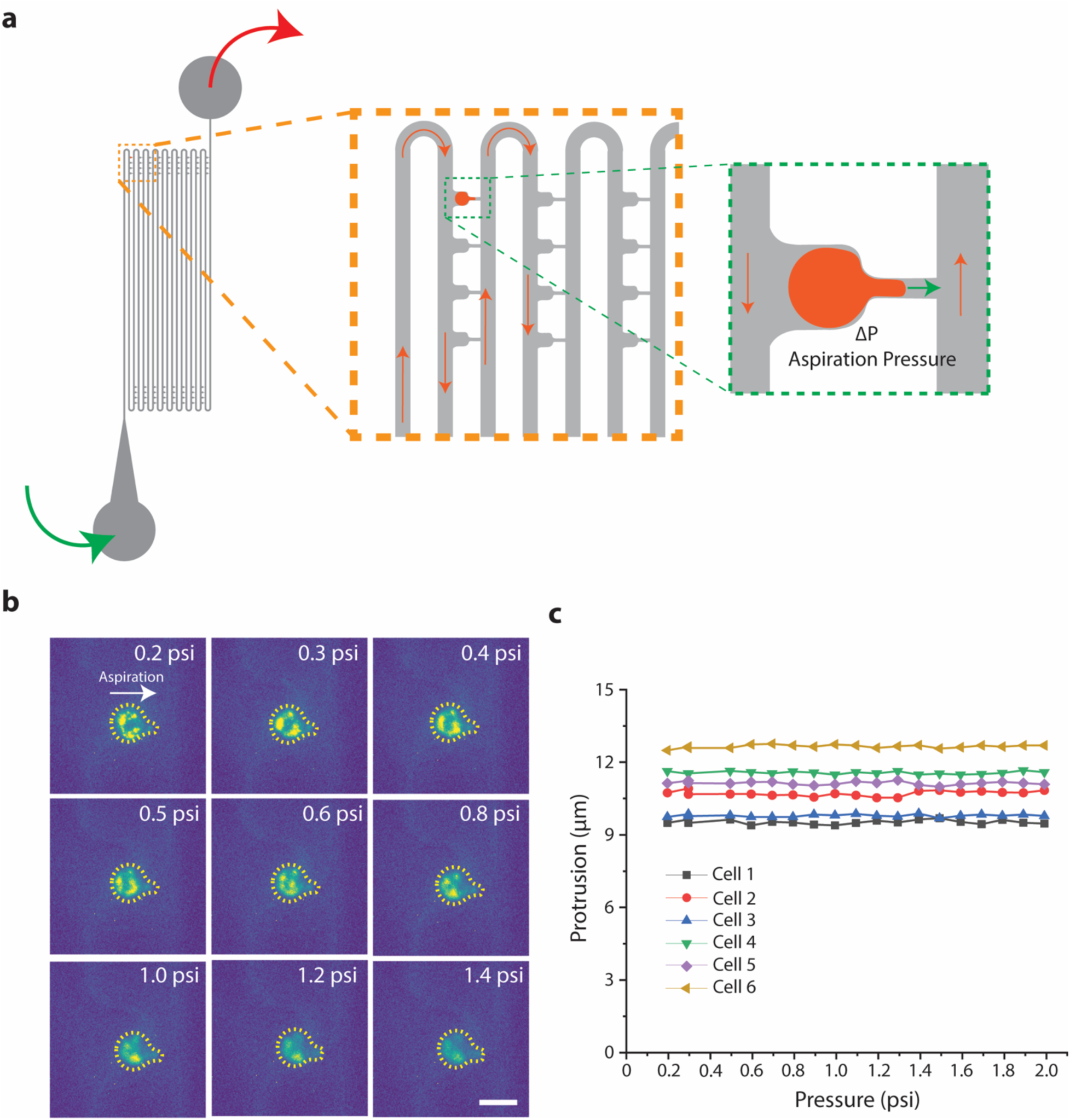
Ground control experiment of microfluidic micropipette aspiration of hFOB cells labeled with Vybrant DiO in the CubeLab. **(a)** Schematic of microfluidic pipette aspiration device indicated direction of fluid flow and direction of aspiration. **(b)** Fluorescent representative images of aspirated hFOB cells. Images show cell protrusion at increasing applied pressure. Scale bar is 20 µm. **(c)** Protrusion profile of 6 cells as a function of applied pressure.

### ISS experiment investigating human osteoblast stiffness under microgravity

We launched the ISS experiment to measure osteoblast stiffness for the first time on NG-18, but the experiment did not yield any results due to issues related to cell capture in our microfluidic chip. The reason for this was an issue with the Northrop Grumman Cygnus vehicle which only deployed one of its two solar arrays after separation from the Antares launch vehicle. This resulted in the payload needing to be shut off en route to the ISS. This power-off resulted in the incubator holding the cells cooling to ∼25 °C for the 28 hours that it was powered off. We suspect this resulted in poor cell viability. We re-launched the experiment on SpX-27 and executed the experiments following the timeline in **Supplemental Table 1**. Prior to the launch, one chip-manifold interface did not fully seal and caused fluid to leak out from the chip. The remaining three chips were successfully loaded without issue. Therefore, on-orbit data was collected from three microfluidic chips. hFOB-seeded culture bags were loaded into the CubeLab approximately 56 hours prior to launch. The Cargo Dragon capsule launched on March 15, 2023 and docked with ISS 11 hours after launch. Approximately 35 hours after launch, the CubeLab was installed on the ISS. Cell lifting and resuspension of hFOB cultures began shortly after installation. Seeding of the first PDMS chip was initiated 40 hours after launch, followed by initiation of the concurrent pressurization and imaging protocols. Several brightfield (2-5 per chip) and fluorescence (2-5 per chip) images were acquired from each chip, capturing all trapped cells at each incremental pressure from 0.2 to 2.0 psi, with 2 minutes at each pressure increment. Cell seeding, pressurization, and imaging of each chip occurred after the previous chip was completed. Imaging data was the primary endpoint for this experiment. Therefore, when the experimental protocol was complete, the payload was powered off and stored on the ISS until return to Earth.

For the ISS experiment, some fluorescence images had background noise which may be attributable to autofluorescence from the PEEK (polyetheretherketone) manifold on which the chips were mounted (Figure 4a). Similar to the ground experiment, we found that the protrusion length into the microfluidic pipette did not change with increasing pressure (Figure 4b). Since we did not see changes to deformation with increasing pressures, we could not determine cell stiffness from these measurements. However, since both experiments had cells subjected to 0.2 psi as the initial pressure, we compared the initial protrusion lengths between ground and ISS samples. The cells on the ISS had a larger protrusion length in the microfluidic pipette channel (Figure 4c), indicating they could be softer. However, we were not able to deduce more from this data alone.

**Figure 4.**
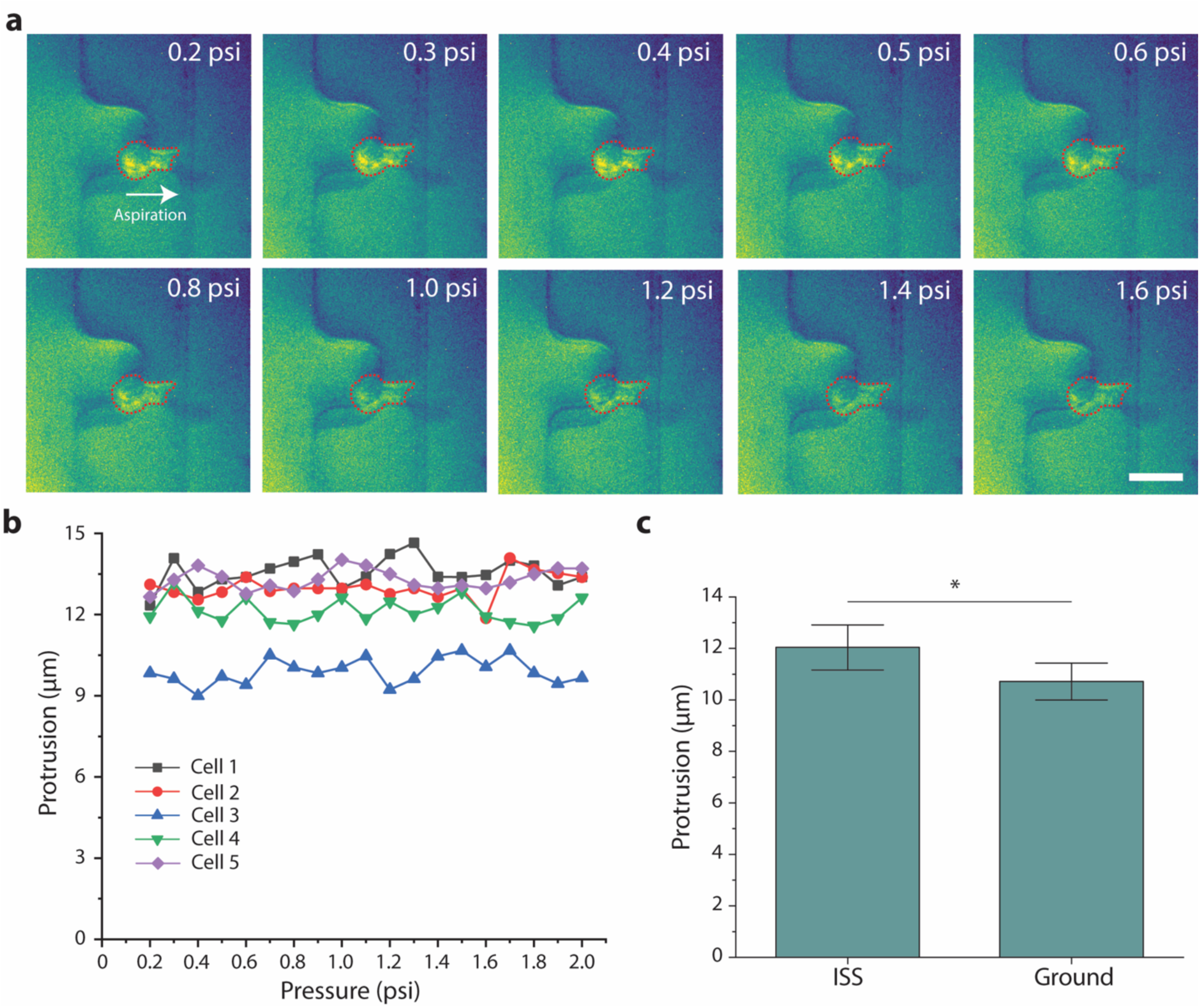
Microfluidic micropipette aspiration of hFOB cells labeled with BioTracker 490 on the ISS. **(a)** Representative images of aspirated fluorescent hFOB cells. Images show cell protrusion at increasing pressure. Scale bar is 20 µm. **(b)** Protrusion profile of 5 cells as a function of pressure. **(c)** Bar graphs comparing protrusion of ISS and control (ground) samples at 0.2 psi. *N*_ISS_ = 6, *N*_Ground_ = 5. Error bars indicate S.E.M..

### Hardware design for automated pressure application to hydrogel-embedded 3D spheroids for the ISS

Our second ISS experiment is aimed at exerting pressure on osteoblast spheroids and testing if such mechanical loading would reverse the effect of microgravity on BMP signaling. Our lab has shown that normal compressive stress on 2D cell monolayer promotes actin network assembly and invasive phenotype of breast cancer cells^25^. Furthermore, we found that compressive stress could accelerate wound closure of breast cancer cells in 2D^24^. These prior studies support the idea that cells regulate cytoskeleton dynamics in response to compressive stress, thereby regulating YAP nuclear translocation. On 2D substrates, It is known that cell spreading correlates with cell tension^22,23^. Using microcontact printed substrates with different cell spreading areas, we found that hFBOs exhibited cell tension-dependent YAP nuclear localization in 2D (**Supplemental Figure 2**).

The microgravity environment is known to induce 3D spheroid formation, and multicellular spheroids embedded in hydrogels have become increasingly popular as these *in vitro* 3D culture models have more physiological relevance compared to traditional 2D cell culture^26,27^. Thus, for this experiment, we tested the effect of compressive stress on BMP signaling using human osteoblast spheroids.

With the same aforementioned space constraint of the CubeLab and the need to have different sample conditions, we opted for a well-plate design format to accommodate 12 inserts with two collagen gels per insert. Figure 5a shows the well-plate layout and the fluidic diagram that allows independent control of fluid handling and pressurization. Figure 5b shows the manufactured well-plate interfaced with the circuit board, and Figure 5c shows the fully assembled CubeLab. Pressurization is applied using a pumping feedback system allowing for compression of fluid to specific pressure setpoints in a closed system. Although it is possible to apply more than 15 psi of pressure without leakage, we found that pressure in excess of 5 psi would damage the collagen hydrogel, potentially damaging the embedded cells. Thus, we decided to use 0 (as control), and nominally 2 and 4 psi for our experiment. Please note this pressure is different from the first experiment as the pressure serves different purposes. The fluidic route is designed such that an adjacent pair of wells would be subjected to the same pressure (i.e., 1 and 2, 3 and 4, 5 and 6 have the same pressure). The hardware can hold pressure well over time (Figure 5d). Since we target a final pressure of 2 and 4 psi, the initial pressure is higher. Nevertheless, the changing pressure is not a major issue for our experiments. Since we are limited by the number of spheroids we can generate from a practical standpoint, another design feature is a custom PDMS insert that constrains the volume of collagen gel to 120 µL. In addition, the insert accommodates two gels per well so that we could have a technical replicate built into the experiment. Finally, the independent valve control allows the perfusion of different solutions. At the end of the experiment, half of the samples were fixed with paraformaldehyde (PFA) for immunofluorescence analysis and half were treated with RNAlater for RNA-Seq analysis.

**Figure 5.**
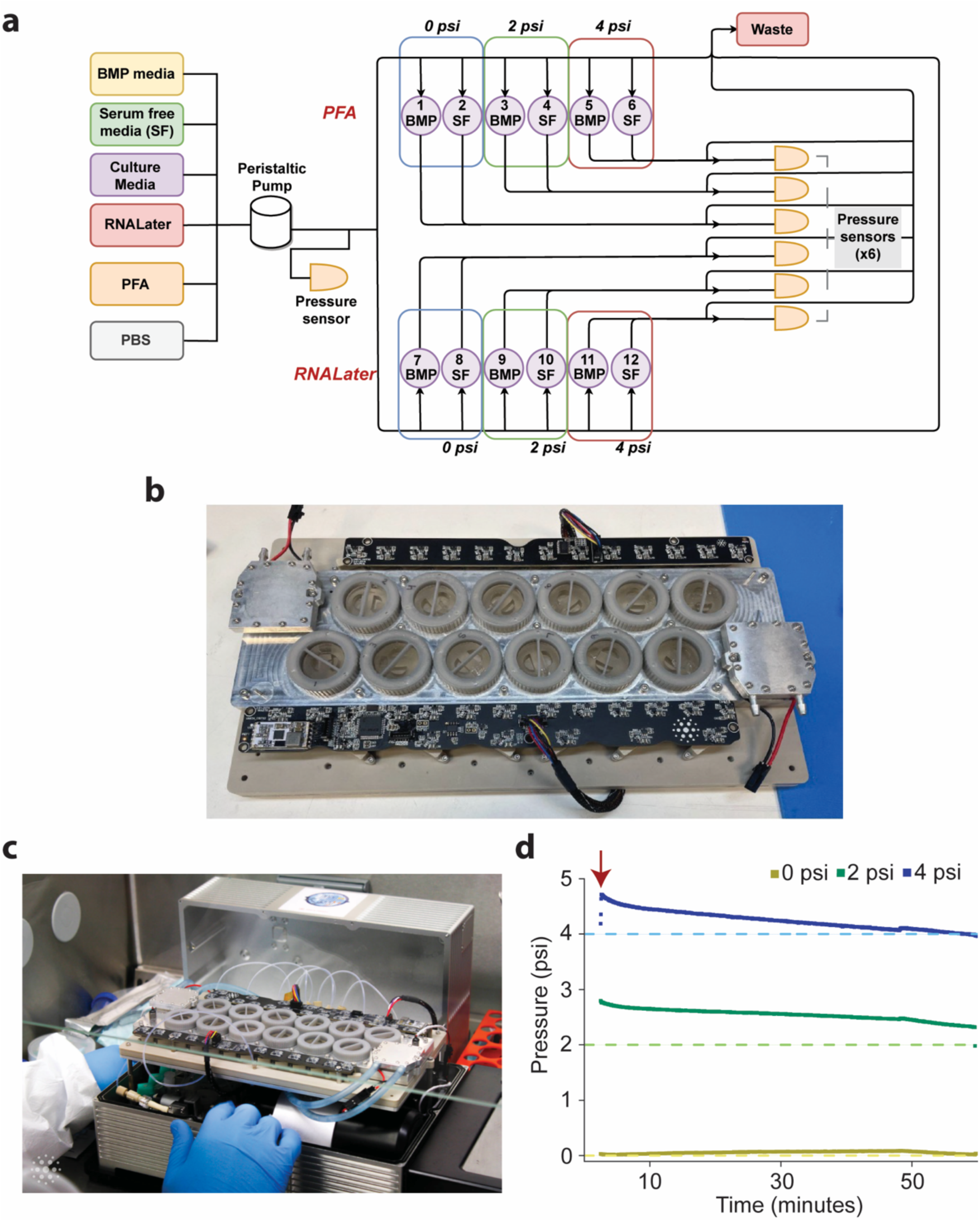
CubeLab for spheroid pressurization experiment. (a) Space Tango’s hardware fluidic diagram illustrates the fluidic routes and components of the integrated CubeLab. (b) Photo of Space Tango’s 12-well plate interfaced with the circuit board. (c) Photo of the fully assembled CubeLab in a biological safety cabinet. (d) Plot of the average of two pressure traces for the 0, 2, and 4 psi wells over the duration of the experiment. Dashed lines show the target pressure level; solid lines show pressure measurements. The red arrow denotes the onset of pressure application.

### Ground testing and experiment of spheroid pressurization with and without BMP2 stimulation

We embedded ∼1000 hFOB spheroids in 3 ml of 3 mg/ml of collagen gel in 12 custom PDMS inserts. Each insert has 2 wells as replicates with each well holding 120 µl of spheroid solution. The spheroids were subjected to 0, 2, and 4 psi for 1 hour with and without 3 hours of pre-treatment of 100 ng/ml of BMP2 stimulation. Half of the samples were washed with PBS followed by PFA fixation for immunofluorescence. The other half of the samples were placed in RNAlater and frozen for RNA extraction and RNA-Seq analysis. The experimental conditions for the different wells are detailed in **Supplemental Table 2.**

The ground experiment using flight hardware followed a timeline that matched the flight schedule where we maintained incubation and monitored data over the first 5-6 days to simulate the launch, ascent to the ISS, and on the ISS. The experiment commenced on day 6 with 12 hours of serum starvation prior to BMP2 stimulation and the pressurization protocol. At the end of the ground experiments, the samples were fixed and moved to cold storage.

The hFOB spheroids were stained for pSMAD1/5/9, YAP, F-actin, and nuclei and imaged by spinning disk fluorescence confocal microscopy (Figure 6a). Actin networks depolymerized in spheroids in conditions with applied pressure, with or without BMP2 treatment. Notably, BMP2 treatment reduced F-actin in ambient conditions as well. The addition of BMP2 increased pSMAD1/5/9 at 2 psi but did not increase pSMAD1/5/9 under ambient and 4 psi conditions (Figure 6b). Since cells are densely packed in a spheroid and F-actin staining is significantly lower in the pressure conditions, discerning nuclear vs. cytoplasmic YAP from our images was not possible. However, it was very apparent that YAP expression was markedly higher with compressive pressure and with the addition of BMP2, suggesting that the increase in YAP in these conditions also corresponds to an increased amount of both nuclear and cytosolic YAP.

**Figure 6.**
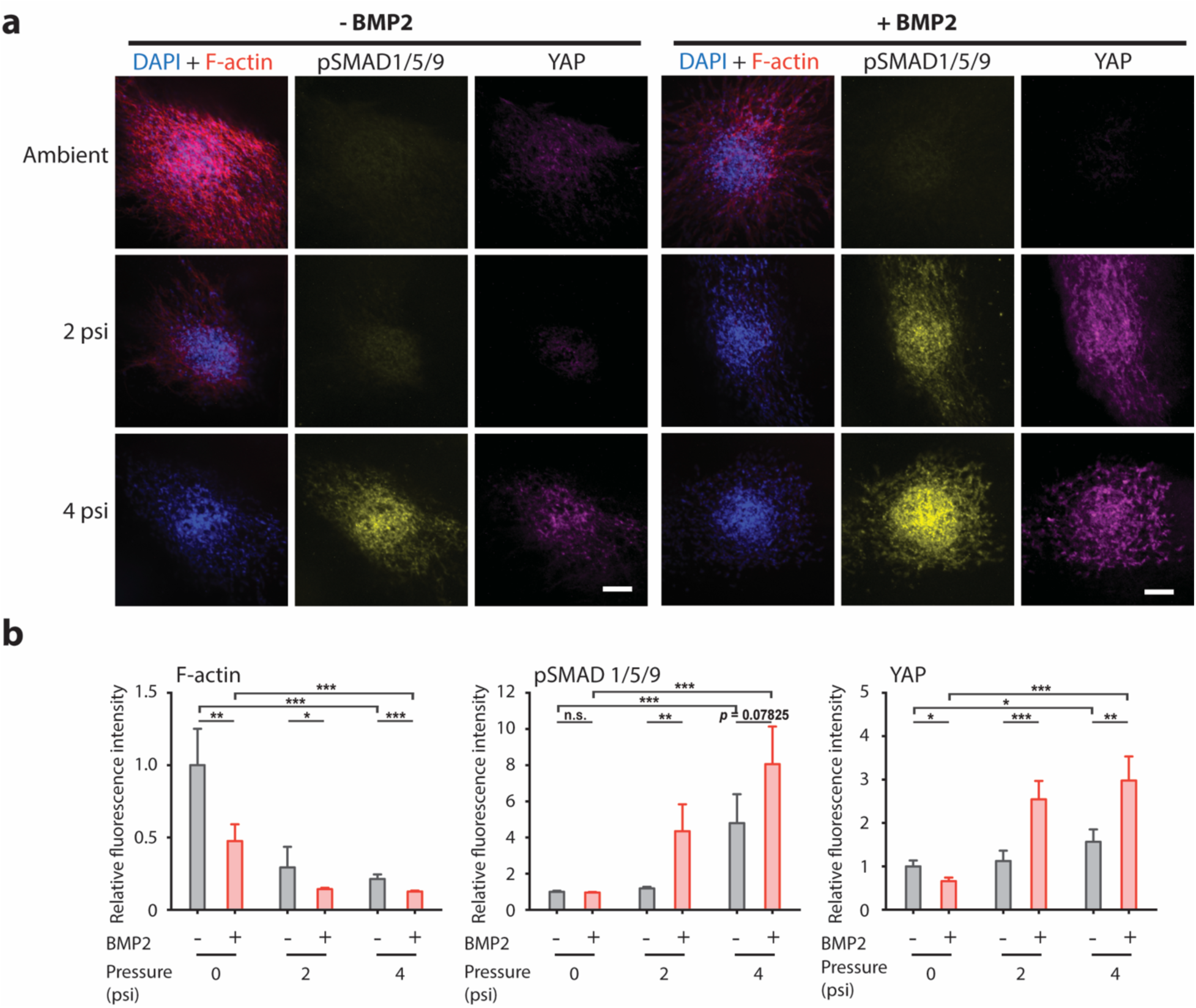
Analysis of hFOB spheroid pressurization experiment on the ground. **(a)** Representative fluorescence images of hFOB spheroids encapsulated in collagen gels. On the ground, the spheroids were pre-treated with 100 ng/ml of BMP2 stimulation for 3 hours (right) and subjected to 1 hour of 0, 2, and 4 psi. Following BMP2 stimulation and application of compressive stress, the samples were frozen and subsequently fixed and immunostained for DAPI, F-actin, pSMAD 1/5/9 and YAP. Scale bar is 100 µm. **(b)** Plots of relative fluorescence intensity of F-actin (left), pSMAD 1/5/9 (middle), and YAP (right) for hFOB spheroids subjected to different levels of compressive stress and with (red) or without (gray) BMP2 stimulation. n = 9–18 spheroids per condition. * = *p* < 0.05, ** = *p* < 0.01, *** = *p* < 0.001.

To further explore how hFOB spheroids react to differences in pressure with BMP2 treatment, gene expression profiles from RNA-sequencing were compared using principal component analysis (PCA)^28^. Spheroids in the 0 and 4 psi conditions were separated along the first principal component regardless of BMP2 treatment while the spheroids subjected to 2 psi were more similar to the 0 psi samples when treated with BMP2 and more similar to the 4 psi samples without BMP2 treatment (Figure 7a). Similarly, gene expression in the Hippo signaling pathway was consistent between BMP2 treatments within the 0 and 4 psi conditions compared to the 2 psi spheroids, which showed changes in expression and downregulation of YAP1 with BMP2 treatment (Figure 7b). To identify changes in transcriptional pathways within each pressure condition as a result of BMP2 treatment, we ranked genes by the probability that they were differentially expressed (**Supplemental Figure 3**) and performed pathway annotation using gene set enrichment analysis^29^. We observed upregulation of pathways related to extracellular matrix organization in response to BMP2 treatment in both the 0 and 4 psi conditions and downregulation of cell-cell communication through gap junctions in 0 psi (Figure 7c). BMP2 treatment activated the Wnt and Hedgehog signaling pathways at 2 psi in addition to nuclear signaling through Nuclear factor-like 2 (NRF2), encoded by the *NFE2L2* gene. Furthermore, glucose metabolism was differentially regulated across conditions, where both glycolysis and gluconeogenesis were upregulated and downregulated in response to BMP2 treatment at 4 psi and 0 psi, respectively. Overall, these results indicate that BMP2 treatment at 2 psi in osteoblast spheroids induces a shift between the distinct transcriptional profiles at 0 and 4 psi, which are characterized by remodeling of the extracellular matrix and differences in glucose metabolism.

**Figure 7:**
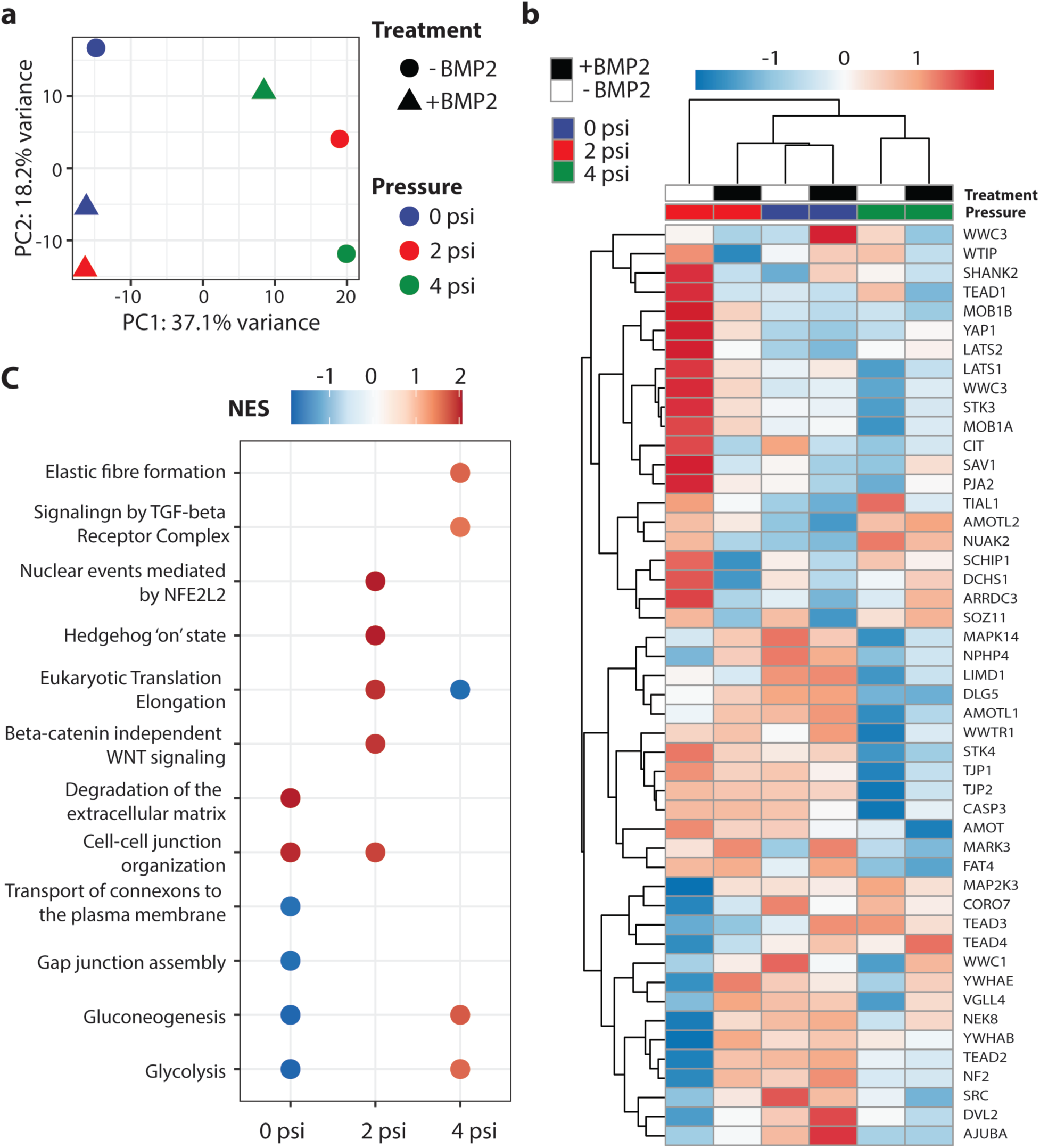
Transcriptomic analysis reveals fluctuations in YAP/TAZ signaling. **(a)** Principal component analysis (PCA) of ground control samples. **(b)** Row-scaled heatmap of gene expression in the Hippo signaling pathway (R-HSA-2028269 from the Reactome database and GO:0035329 from the GO database). **(c)** Significant Reactome terms (p_adj_ < 0.05) from gene set enrichment analysis (GSEA) of ranked differences in expression between BMP2-treated and untreated samples within each pressure group. Points are colored by normalized enrichment score (NES) with positive scores indicating upregulation in the BMP2-treated sample

### ISS experiment investigating the effect of BMP stimulation on human osteoblast spheroids in microgravity

The ISS experiment was launched on NG-18 and the experiment ran smoothly once installed on the ISS (**Supplemental Table 3**). hFOB spheroids were loaded into the CubeLab approximately 66 hours prior to launch. The Cygnus capsule launched on November 7, 2022 and docked with ISS 51 hours later. Serum starvation of spheroid cultures began approximately 12 hours after the CubeLab was installed on the ISS. Treatment with or without BMP was initiated after the 12-hour serum starvation period was complete with the pressurization routine starting 2 hours later. The pressurization protocol also operated without any issues, as supported by the pressure, temperature, and flow sensor readings throughout the experiment. The samples were fixed and subsequently frozen at the end of the experiment and returned to Earth with SpX-26 on January 11, 2023.

Although microgravity is known to disrupt the actin cytoskeleton, we found that the application of 2 and 4 psi on hFOB spheroids significantly reduced F-actin, similar to our ground experiment, irrespective of BMP2 simulation (**Figure 8a, b**). BMP2 addition in the absence of applied pressure also significantly reduced F-actin. Interestingly, pSMAD1/5/9 was also elevated in pressurized samples. There was a slight increase in pSMAD1/5/9 when BMP2 was added to the samples pressurized by 4 psi. These results support the idea that the application of external pressure would promote BMP2-stimulated SMAD signaling. However, it is worth noting that SMAD1/5/9 was phosphorylated even in the absence of BMP2 simulation when pressure was applied. YAP expression was comparatively low in all the conditions except at 4 psi with BMP2 addition. The results from the ISS are in general agreement with the ground experiment. For all conditions, hFOB spheroids in microgravity exhibited significantly lower F-actin compared to the ground samples. Both YAP expression and BMP2-stimulated pSMAD signaling were promoted by the application of pressure. We found that microgravity led to greater actin disassembly and lowered YAP expression (Figure 8c). Our result suggests that compression can indeed restore YAP activity in microgravity.

**Figure 8.**
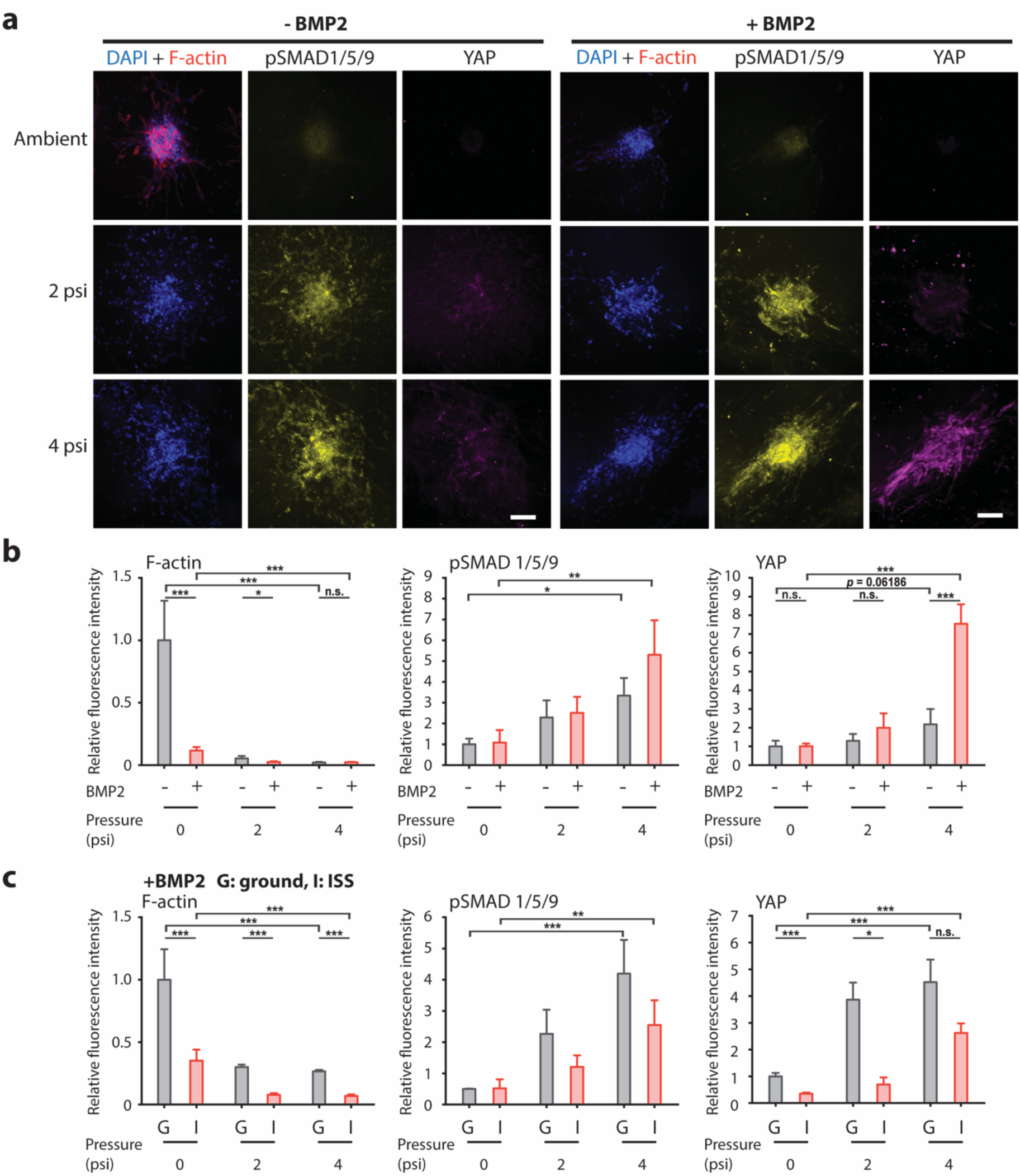
Analysis of hFOB spheroid pressurization experiment on the ISS. **(a)** Representative fluorescence images of hFOB spheroids encapsulated in collagen gels. On the ISS, the spheroids were pre-treated with 100 ng/ml of BMP2 stimulation for 3 hours (right) and subjected to 1 hour of 0, 2, and 4 psi. The samples were fixed and stored at -20 °C on the ISS and immunostained for DAPI, F-actin, pSMAD 1/5/9 and YAP on the ground. Scale bar is 100 µm. **(b)** Plots of relative fluorescence intensity of F-actin (left), pSMAD 1/5/9 (middle), and YAP (right) for hFOB spheroids subjected to different levels of compressive stress and with (red) or without (gray) BMP2 stimulation. **(c)** Plots of relative fluorescence intensity for BMP2-stimulated hFOB spheroids subjected to 1 hour of 0, 2, and 4 psi on the ground (gray) and on the ISS (red). n = 6–16 spheroids per condition. * = *p* < 0.05, ** = *p* < 0.01, *** = *p* < 0.001.

Although we had successfully obtained RNA samples from our ground control experiment using flight hardware, unfortunately, we did not obtain RNA samples of sufficient quality with the flight samples (RNA integrity number of ∼ 2.5 with 10 being the highest quality). The low-quality RNA prevented us from performing RNA-Seq or RT-PCR analysis. We suspect RNA degradation occurred as a result of multiple freeze-thaw cycles during the journey from the ISS to processing RNA samples in the lab. When processing the RNAlater samples with Trizol, we noticed that the gels were difficult to dissolve, and we were unable to recover a sufficient number of intact spheroids from the gels.

## DISCUSSION

In this study, we aimed to explore the intersection of BMP/YAP signaling in human osteoblasts in the context of microgravity. Flight hardware was developed for two independent ISS experiments by Space Tango. Subsystems went through iterative designs and extensive ground testing. For human osteoblast stiffness measurement, we find that osteoblast deformation did not vary as a function of applied pressure on the ISS. This is unexpected and the exact reason for this behavior is unclear. Since actin is depolymerized in microgravity, it is possible that the bulky nucleus is stuck at the micropipette and the softer cytoplasm is fully aspirated at the lowest applied pressure. We previously found human osteoblasts exhibit a more disorganized actin cytoskeleton and lower stiffness in a simulated microgravity condition^19^. Thus, our ISS and ground experiments are consistent with our previous findings.

For the osteoblast spheroid pressurization experiments, our data indicated microgravity impacts F-actin networks but did not impact BMP2-stimulated pSMAD1/5/9. YAP expression in osteoblasts was negatively impacted by microgravity. Both pSMAD1/5/9 and YAP levels were significantly higher in the pressurized samples. At 2 and 4 psi, spheroids in microgravity exhibited lower YAP and pSMAD1/5/9 expression compared to spheroids in the ground experiment. Our RNA-Seq datasets further substantiated a decrease in YAP1 with BMP2 treatment at 2 psi and differences in extracellular matrix remodeling. In a previous study, it was found that the YAP pathway is differentially expressed in blood cells during space flight^2^. It is possible that cell viability might have been influenced at 4 psi, accounting for the lack of increased pSMAD with BMP2 treatment.

Nabavi *et al.* discovered that when osteoblasts are subjected to 5 days of microgravity, their cytoskeleton becomes disrupted, leading to disassembled F-actin fibers and shorter, wavier microtubules^30^. Previous research has demonstrated that interfering with F-actin or inhibiting myosin contraction abrogates YAP/TAZ mechanotransduction^10^. These findings strongly indicate that when the integrity of the actin cytoskeleton is compromised, YAP/TAZ mechanotransduction is adversely impacted. Furthermore, YAP is a well-known key effector molecule for Hippo signaling that controls organ size by modulating cell proliferation and apoptosis. It is well established that the actin cytoskeleton regulates Hippo signaling ^31,32^ and therefore YAP activity via kinases LATS1/2. An increase in F-actin led to YAP de-phosphorylation and nuclear localization in HeLa cells^32^. Our finding of reduced F-actin at 2 and 4 psi yet increased YAP expression appears to contradict this. How pressure regulates F-actin disassembly in our experiment is not immediately clear. However, it is also known that YAP can be activated in a Hippo-independent manner^33,34^. Interestingly, YAP nuclear translocation can be induced by the application of force, even when actin is depolymerized^35^. It is thought that force increases YAP nuclear import by reducing mechanical restriction in nuclear pores. The increased YAP intensity in the pressure conditions could be due to this YAP forced-entry mechanism, which correlates with the increase in glycolysis^36^ we observed in our RNA-Seq datasets. Perhaps more interesting is the finding that pressure application led to transcriptional upregulation of YAP. To our knowledge, this has not been observed previously and will require more studies.

It is important to note the limitations of our study. It is inherently difficult to carry out experiments on the ISS due to space constraints. Thus, the scope of our investigation is limited, and we were constrained by conducting each of the ISS experiments only once via one flight per experiment. Numerous factors were considered during experimental design to ensure that we had all the conditions covered and that we had sufficient samples to analyze. The space constraint and the automated imaging requirement limited the number of microfluidic chips to four. However, limiting the number of chips to four does not intrinsically compromise the number of data points for statistical analysis since a single chip, in an ideal experimental condition, is capable of trapping 64 cells. Automating cell trypsinization and resuspension is a unique challenge terrestrially and likewise was a challenge within the CubeLab. Our flight provided an opportunity to develop a hardware solution using a vibrating disc motor for cell resuspension which may prove to be useful for other future flights with similar requirements. The brightfield and fluorescence imaging in the CubeLab are very different from an inverted fluorescence microscope in a research lab. These differences stem primarily from the compact size and full automation of the system. The image quality with out-of-focus and autofluorescence from PEEK posed some challenges to image analysis, making it difficult to draw firm conclusions. For the second experiment, the design of our pressure well plate setup proved to be robust for sustaining pressure application to biological specimens. Although we successfully carried out immunofluorescence experiments from the returned frozen spheroid samples, the multiple freeze-thaw cycles likely contributed to RNA degradation, so we could not proceed with RNA-Seq or RT-PCR analysis. Finally, we did not include cell viability markers in our experiments. However, the cell stiffness measurement itself is a short experiment that we do not expect to impact cell viability, but the automated fluidic setup lengthens the experimental protocol compared to what is done on Earth. For the spheroid pressurization experiment, it is possible the pressure could impact cell viability, as noted earlier. However, our spheroid system was tested on Earth for over 10 days without a viability issue with regular media changes.

While manual experiments on Earth can be continuously improved and human interactions with the experiments ensure any issues can be quickly remedied, conducting automated experiments on the ISS has its own challenges. For example, changing clogged chips, high-pressure flushing for cleaning, and hands-on experimental reboot to eliminate residual pressure or trapped air bubbles cannot be easily resolved in space and are more quickly addressed when performing experiments manually on the ground. Hardware development includes multiple rounds of subsystem Engineering Verification Tests (EVTs) that provide opportunities to test and validate hardware systems. Automated space-based hardware provides the same advantages seen with terrestrial hardware, and improves repeatability and reliability from experiment to experiment, thereby increasing the quality of results. On the ISS, automated hardware reduces the requirement of crew time, a significant factor in cost and schedule for flight opportunities. Considering the complex nature of performing automated cell biological experiments in space, the outcomes of the experiments were satisfactory.

In summary, our work opens the door for future cellular biology studies on the ISS or in space. We successfully implemented two experiments with automated instrumentation. We anticipate the innovation of hardware and documented protocols can aid future studies on the ISS. Although the findings of our study require additional research to validate, they nevertheless provide the first data on pSMAD and YAP in human osteoblasts in microgravity conditions and how pressure might restore YAP activity. These findings have implications for human health benefits on Earth in understanding and preventing osteoporosis. Given that bone loss is among the pronounced problems astronauts face as a result of an extended stay in space requiring preservation approaches such as resistance exercises in an attempt to rescue bone loss, our study also provides ideas for promoting bone health in future long-term space travel.

## METHODS

### Cell culture and spheroid generation

hFOB 1.19 cells (ATCC CRL-3602) were cultured in 1:1 DMEM/F12 medium (Gibco) supplemented with 0.3 mg/ml G418 and 10% FBS. Cells were cultured in a humidified atmosphere containing 5% CO_2_ at 33.5°C.

For mechanical characterization of single cells, hFOB cells were stained using either BioTracker 490 Green (Thermo Fisher) cytoplasmic membrane dye or Vybrant DiO Cell-Labeling Solution (Invitrogen). 5 µM of BioTracker 490 or 5 µL of Vybrant DiO per 1 ml media was added to 1x10^6^ cells/ml suspended in media and incubated for 20 minutes followed by 3 subsequent washes with PBS. The two membrane dyes have comparable spectral properties and can both maintain cell membrane staining for over a week. 10 ml of cell solution was then transferred to thin-walled, gas exchange Kiyatec bags at 50,000 cells/ml and integrated into the CubeLab at science loading. Four culture bags were loaded into the CubeLab. One media exchange was performed, during ascent to the ISS, by exchanging 50% of the spent media with fresh culture media.

For spheroid generation, 3D spheroids were formed using custom plates, which consist of microfabricated inverted pyramidal microwells and work by adding single-cell suspension to the wells containing microwells and centrifugation distributes cells evenly into the microwells. ∼1,000 hFOBs per microwell (200 microwells per well on a 96-well plate) were spun down at 300g for 5 min. Spheroids were allowed to form overnight. After spheroid formation, spheroids were encapsulated in collagen hydrogel consisting of 3 mg/mL rat-tail collagen (Advanced BioMatrix). The encapsulated spheroids were cultured in custom inserts fabricated by Space Tango, in a humidified environment containing 5% CO_2_ at 33.5°C and submerged in hFOB culture media. At the time of science loading, custom inserts containing cultured, hydrogel-encapsulated spheroids were loaded into Space Tango’s CubeLab 12-well system. For BMP2 stimulation, the samples were serum-starved for 12 h and then treated with 100 ng/ml BMP2 (R&D Systems) for 3 h. Following serum starvation, half of the wells received BMP2, and the other half received serum-free media only. Hydraulically applied pressure was applied to the encapsulated spheroids at either 0 psi, 2 psi, or 4 psi, and pressurization lasted for 1 hour each. Following pressurization, all samples were washed with Phosphate-Buffered Saline (PBS) (Gibco) before fixation. Half of the wells were fixed with 4% paraformaldehyde (Thermo Scientific) and stored in PBS and the other half were fixed and stored in RNAlater (Invitrogen). The entire CubeLab, containing all fixed samples, was stored at -20°C until its return to Earth.

### Flight hardware

Space Tango flight and ground hardware were designed to support hFOB 2D and 3D culture maintenance, fluid conveyance and routing, thermal control, pressurization, and imaging. The wetted materials selected for use within the CubeLab were certified as biocompatible and sterilizable. The wetted fluidic system was sterilized, assembled in a sterile manner, and subsequently tested and confirmed to be sterile prior to loading experimental fluids and biology. Fluids requiring cold storage were maintained in a thermally controlled environment at 4°C ± 3°C. hFOB cultures were maintained at 33.5°C ± 3°C and 5% CO_2_ ± 1% (passive maintenance).

A key feature of Space Tango’s automated hardware is its ability to move fluids in a sterile manner with precision and reliability. This is accomplished using machined fluidic manifolds made from biocompatible engineered plastics. These manifolds were custom-designed with fluid routing cross-section profiles optimized for bubble mitigation in microgravity as well as independent fluidic routing through valving specific to each science interface system used (i.e., PDMS chips for hFOBs mechanical measurement or a well plate for spheroid pressurization).

A key requirement for each experiment was pressurization. Pressurization of the system was accomplished in the same way for both experimental systems, PDMS chip and well plate. Each fluidic circuit included a programmable syringe in line with the manifold to hydraulically achieve the required pressures for each experiment. This included a feedback loop that pulled measurements at various points within the fluid manifold to ensure the required pressures were met and maintained once the valving was closed.

### Microfluidic micropipette aspiration experiments

hFOB cells grown in 2D were transferred from their culture bag (Kiyatec) to the PDMS chips using an innovative design solution for cell resuspension. The protocol for hFOB detachment from the culture bags was adapted from the TrypLE Express manufacturer’s protocol. Briefly, hFOB culture media was removed from the bag, 2mL of TrypLE Express (Gibco) was added to the cell monolayer, and a vibrating disc motor was used to agitate the monolayer. The monolayer was incubated at 37°C for 10 minutes followed by an additional round of agitation for cell lifting. Cell culture media (8 mL) was added to the cell suspension to create the optimal cell density required for seeding the PDMS chips.

To mechanically characterize hFOB cells, a multilayer PDMS-based microfluidic pipette array was used^16^. Designed to be highly robust and rapid, the working principle of our microfluidic chip is similar to the glass capillary-based micropipette aspiration method. A serpentine main channel is connected by a cell-trapping chamber and a micropipette channel. Due to laminar friction throughout the main channel and head loss at turns, a pressure difference will trap and exert a suction force onto single cells. There are 64 trapping/aspirating chambers throughout the device. Four microfluidic chips were integrated into the CubeLab to automate the mechanical characterization of hFOB cells. Fluorescently labeled cells, using BioTracker 490 or Vybrant DiO, were loaded into cell trapping structures at 0.2 psi and aspirated by increasing the pressure in increments of 0.1 psi in the range of 0.2-2 psi resulting in the protrusion of cells into the micropipette channel. Images capturing at least 12 cups, both fluorescence and brightfield, at each pressure increment, were captured using a 20X objective. To avoid time-dependent changes in protrusion due to the viscoelastic property of cells, we waited 2 minutes before capturing images at each pressure increment to allow equilibration of deformation. From the collected images, we measured protrusion lengths at each pressure increment using ImageJ.

### Confocal microscopy

Images were taken using an UplanFL N 10 x/1.30 NA (Olympus) objective on an inverted microscope (Olympus IX-81) equipped with an iXON3 EMCCD camera (Andor Technology), National Instrument DAQ-MX controlled laser (Solamere Technology), and a Yokogawa CSU-X1 spinning disk confocal unit. Z-stack images of spheroids fluorescently labeled for DAPI, YAP, pSMAD and F-actin were taken at excitation wavelengths of 405, 488, 561, and 640 nm, respectively, with a 3 µm z-step.

### Immunofluorescence staining and analysis

hFOB spheroids embedded in collagen gels were washed with PBS and fixed with 4% paraformaldehyde for 1 h, washed with PBS, and permeabilized with 0.1% Triton X-100 in PBS overnight at 4°C. Afterward, the samples were blocked with 3% BSA in PBS overnight at 4°C and then incubated with a rabbit phospho-SMAD1/5/9 antibody at 1:400 (D5B10, Cell Signaling) and a mouse anti-YAP antibody at 1:200 (63.7, Santa Cruz Biotech) in 3% BSA overnight at 4°C. Next, the samples were washed 3× with PBS for 30 minutes per wash and incubated with DAPI, Acti-stain 670 phalloidin, and secondary antibodies in 3% BSA overnight at 4°C. The samples were washed 3× with PBS as described above and imaged by spinning disk confocal microscopy.

For image analysis of experiments nuclear/cytoplasmic YAP ratio, background subtraction was performed in ImageJ, and images were subsequently segmented with MATLAB to generate masks for actin and DAPI channels that were used to measure the mean nuclear and cytoplasmic intensity of each region. To quantify YAP, the total nuclear or cytosolic intensity was calculated, and nuclear intensity was averaged over cytosolic intensity to generate a nuclear/cytoplasmic YAP ratio.

### RNA library preparation

Osteoblast spheroids from the ground experiments were collected in RNAlater and frozen at -80°C for RNA collection. Total RNA was isolated with the miRNeasy Micro kit (Qiagen) following manufacturer instructions. Libraries were prepared using the SMART-seq mRNA kit^28,38^ (Takara Bio) and sequenced on NovaSeq instrument (Illumina) using 151bp paired-end reads. The reads were demultiplexed into FASTQ files using the BCL Convert Conversion Software (Illumina, v4) and the adapters were trimmed using Cutadapt v2.3 ^39^. The reads were then mapped to the human GRCh38 reference genome (ENSEMBL 109) following ENCODE standards using STAR v2.7.8a ^40^ and counts were assigned to genes using RSEM v1.3.3. The final libraries contained an average of 18,095,188 uniquely mapped reads.

### RNA-Seq analysis

The excepted counts from RSEM were filtered to genes with at least 10 counts in at least one sample and transformed to log-scaled counts per million using the edgeR package (v3.40.2) ^41^. Principal component analysis was performed with the top 500 variable genes from the log-scaled transformed counts^42^. To estimate differential expression between conditions without biological replicates, the NOISeq package (v2.42.0) ^43^ was used to simulate 5 technical replicates per sample, each containing 20% of the original sample’s reads with 0.02 variability. The technical replicates were used to estimate for which genes the log2 ratio (M) and difference (D) between counts in each condition was greater than expected compared to a noise distribution. A summary statistic of M and D values (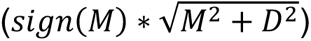) was used to rank genes for gene set enrichment analysis (GSEA) with the Reactome database ^44^ using the fgsea package (v1.24.0) ^45^. A threshold of p_adj_ < 0.05 was used to identify significant Reactome terms.

### Microcontact printing of fibronectin adhesive areas

Micropatterned substrates were made with standard soft lithography techniques to create a silicon master mold^23^. PDMS was prepared by mixing Sylgard-184 elastomer and its accompanying curing agent (Dow Corning) in a 10:1 (w/w) ratio respectively. After degassing, the PDMS mixture was added over the master and cured overnight at 60 °C. Human fibronectin (Invitrogen) was added to stamps at a concentration of 40 µg/ml for 1 hour. Non-adsorbed fibronectin solution was removed by blow drying with air. To prepare the substrate, coverslips were spin-coated with a 9:1 PDMS:hexane solution at a speed of 5,000 rpm for 2 minutes and left to set overnight at room temperature. The coverslips were treated with UV-ozone immediately before placing the stamps in contact with the coated coverslip by briefly applying uniform pressure and then passivated with 0.1% Pluronic F-127 (Sigma-Aldrich) for 1 hour. Stamped substrates were stored at 4 °C in 1x PBS for up to one week.

### Statistical analysis

For protrusion length comparison, a two two-tailed t-test was used with *p* = 0.05 as the significance level. Statistical analysis was carried out in Origin and performed with one-way ANOVA followed by Tukey post-hoc multiple comparisons test. Statistical significance was denoted by asterisks in the figure panels, with * = *p* < 0.05, ** = *p* < 0.01, *** = *p* < 0.001.

## ACKNOWLEDGEMENTS

We thank Annaliza Torres for designing and fabricating the well inserts for the pressurization experiment. N.H.W. acknowledges support from an NIH supplement (EB030031). We acknowledge support from the Bioinformatics Core of the University of Michigan Medical School’s Biomedical Research Core Facilities (RRID:SCR_019168). A.P.L. acknowledges support from the National Science Foundation (1927803) and the Center for the Advancement of Science in Space (GA-2023-9002). T.C. acknowledges support from the Center for the Advancement of Science in Space and the National Aeronautics and Space Administration.

## AUTHOR CONTRIBUTIONS

N.H.W., G.C., T.C., and A.P.L. contributed to the conception of the experiments. N.H.W., G.C., S.J.C., and S.-S.L. contributed to the ground and ISS experiment experimental design. M.S., M.R., D.M., M.H., and P.V. contributed to hardware development, validation, and deployment. P.D. performed RNA library preparation, B.Y. and C.A. performed RNA-Seq analysis. E.A.-M. and Y.Q. conducted ground experiments with osteoblasts. N.H.W. and G.C. analyzed the data from ground and ISS experiments. N.H.W., G.C., S.J.C., B.Y., S.G., T.C., and A.P.L. wrote the manuscript draft. All authors contributed to editing and providing feedback. N.H.W and G.C. contributed equally to this work. T.C. and A.P.L. supervised the project.

## DATA AVAILABILITY STATEMENT

All data generated during this study are included in the manuscript and its supplemental information. Raw data for RNA-Seq are available at the Gene Expression Omnibus (GEO) database (accession number GSE245992). Raw images are available upon request.

