## Supplemental materials for "Cellular mechanotransduction of human osteoblasts in microgravity"

**Supplemental Figures**

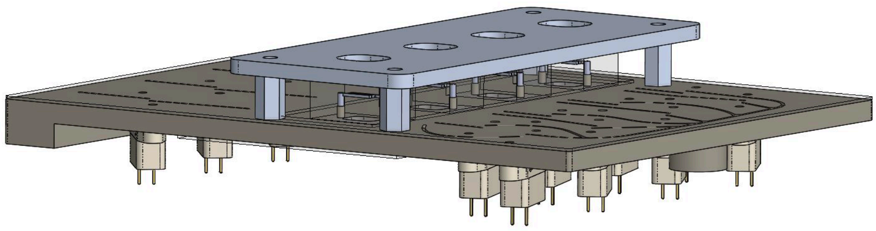

**Supplemental Figure 1.** CAD drawing of the fluidic manifold with a compression plate for holding down the microfluidic chips.

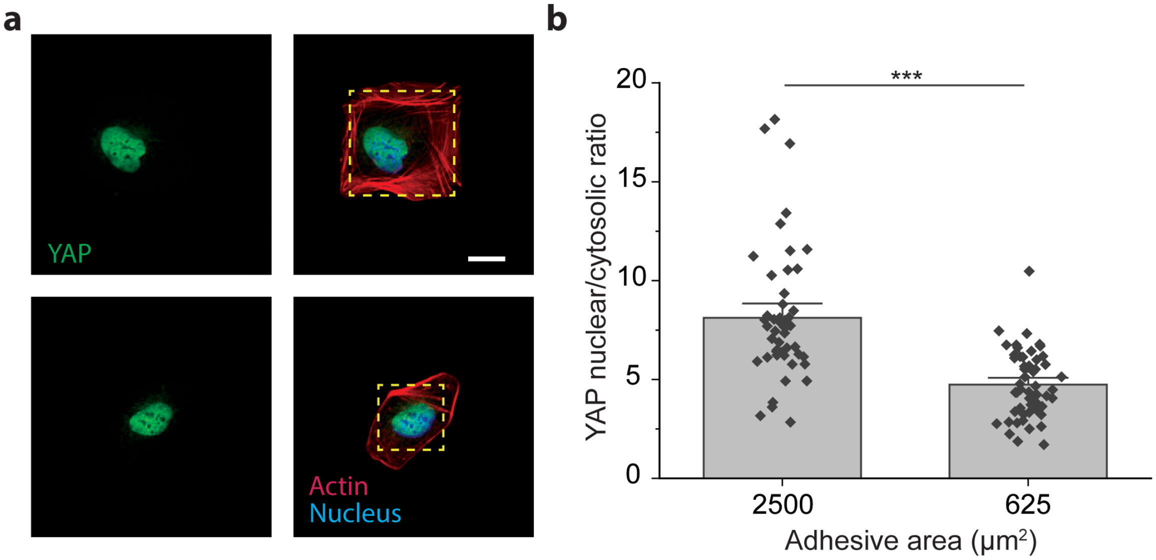


**Supplemental Figure 2. YAP translocates to cytosol at low cell tension. A)** Representative immunofluorescence images of human osteoblasts (hFOBs), on large (2500 $\mu m^2$) and small (625 $\mu m^2$) micropatterns, stained for F-actin, DNA, and YAP. Scale bar is 20 µm. **B)** Quantification of YAP nuclear/cytoplasmic ratio of individual hFOBs from three independent experiments. Error bars denote standard error of the means. *** $p < 0.001$.

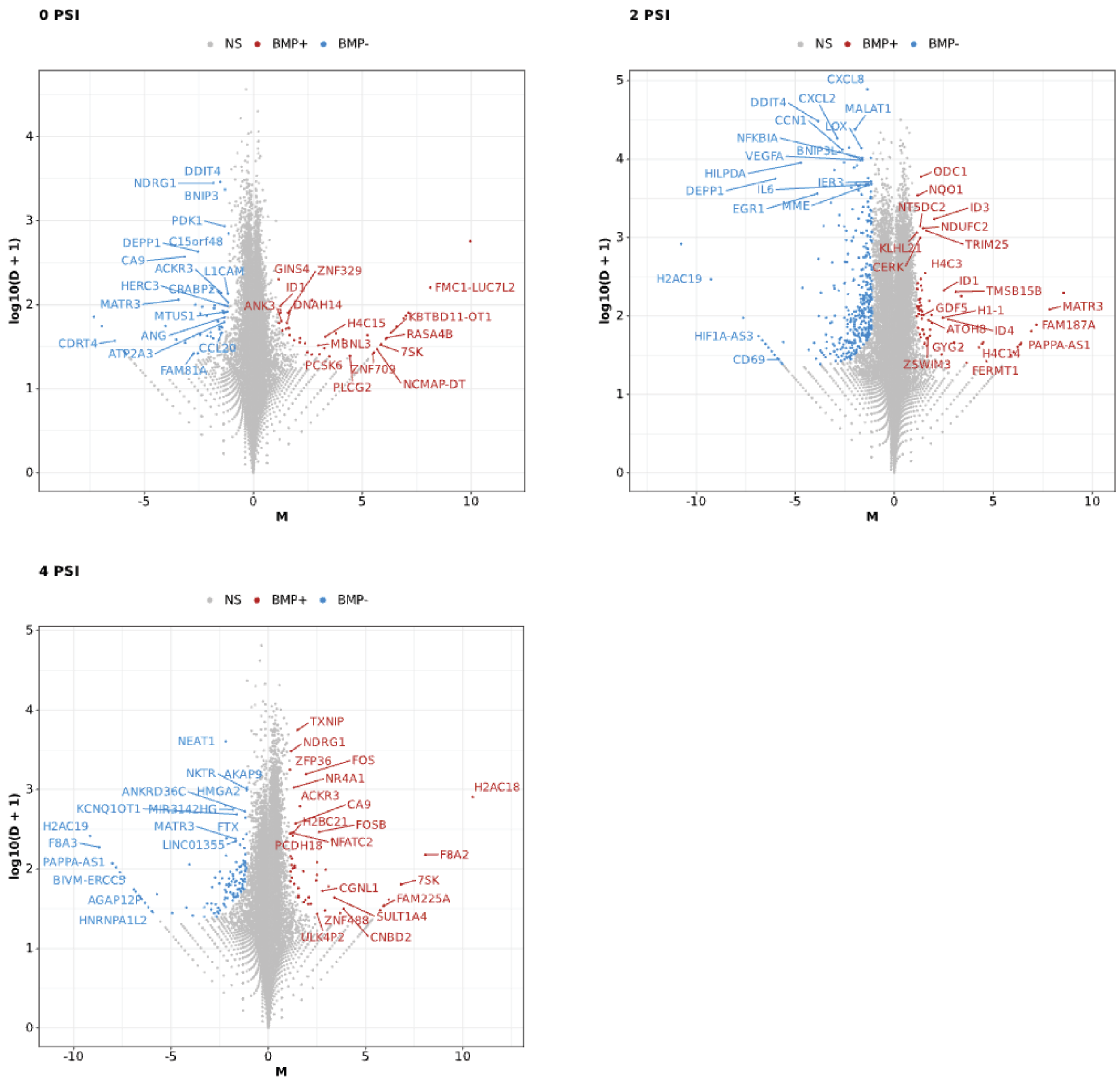


**Supplemental Figure 3. Summary M-D plots showing differences in expression between BMP2 treated and untreated samples within each psi group.** M values indicate log2 ratios between conditions. D values indicate the difference in counts between conditions. Significant genes are identified as those that are >90% likely to be differentially expressed compared to a null distribution.

### Supplemental Tables

**Supplemental Table 1.** Timeline of the ISS experiment on microfluidic aspiration of human osteoblast (flown on SpX-27)

| Date (2023) | Mission time days | Time from loading (hours) | Event (EST) |
|---|---|---|---|
| March 13 | 0 | | 1:00 pm: final loading into CubeLab |
| March 14 | 1 | 28.5 | 5:30 pm: Handover to NASA |
| March 15 | 2 | 55.5 | 8:30 pm: Launch |
| March 16 | 3 | 66.5 | 7:30 am: Dock with ISS |
| March 17 | 4 | 90.5 | 7:50 am: Installed on ISS |
| March 17 | 4 | 95 | 10:30 am: Chip 2 seeding, pressurization, imaging/video |
| March 17 | 4 | 97 | 12:50 pm: Chip 3 seeding, pressurization, imaging/video |
| March 18 | 5 | 124.5 | 4:25 pm: Chip 4 seeding, pressurization, imaging/video |
| Payload remained powered to perform potential additional activities | | | |
| March 24 | 11 | 149 | 5:30 pm: Payload turned off |

**Supplemental Table 2.** Well plate layout for the osteoblast spheroids pressurization experiment.

| Well | 1 | 2 | 3 | 4 | 5 | 6 | 7 | 8 | 9 | 10 | 11 | 12 |
|---|---|---|---|---|---|---|---|---|---|---|---|---|
| Pressure (psi) | 0 | 0 | 2 | 2 | 4 | 4 | 0 | 0 | 2 | 2 | 4 | 4 |
| Ligand condition | BMP | None | BMP | None | BMP | None | BMP | None | BMP | None | BMP | None |
| Fixative | PFA | PFA | PFA | PFA | PFA | PFA | RNA Later | RNA Later | RNA Later | RNA Later | RNA Later | RNA Later |

**Supplemental Table 3.** Timeline of the ISS experiment on osteoblast spheroid pressurization (flown on NG-18)

| Date (2022) | Mission time days | Time from loading (hours) | Event (EST) |
|---|---|---|---|
| November 4 | 0 | | 12:00 pm: Final loading into CubeLab |
| November 5 | 1 | 17 | 5:00 am: Handover to NASA |
| November 6 | 2 | 41 | 5:05 am: Launch attempt 1 |
| November 7 | 3 | 66 | 5:30 am: Launch |
| November 8 | 4 | 95 | 11:18 am: Power shut off due to Cygnus solar panel malfunction |
| November 9 | 5 | 117 | 9:00 am Docked with ISS |
| November 9 | 5 | 123 | 3:30 pm: Power turned back on |
| November 10 | 6 | 139<br>151 | 7:04 am: Installed on to ISS<br>7:00 pm: Start serum starvation |
| November 11 | 7 | 162<br>164<br>165<br>168 | 5:45 am: Start BMP treatment<br>7:36 am: Start pressurization<br>8:45 am: Fixation start<br>11:33 am: Moved to cold stowage |
